# High throughput *in situ* imaging reveals complex ecological behaviour of giant marine mixotrophic protists

**DOI:** 10.1101/2025.08.12.669610

**Authors:** Thelma Panaïotis, Tristan Biard, Louis Carray--Counil, Robin Faillettaz, Jessica Y Luo, Cedric M Guigand, Robert K Cowen, Jean-Olivier Irisson

**Author notes:** **Corresponding author** Thelma Panaïotis.

## Abstract

Mixotrophic organisms have the ability to combine photosynthetic and heterotrophic nutrition, and are ubiquitous in marine ecosystems. Here, we focus on Rhizaria, a diverse group of planktonic protists that included both mixotrophic and non-mixotrophic taxa, and that are particularly delicate and poorly sampled by standard plankton nets. Using high-frequency *in situ* imaging, we study the ecology of these organisms directly in their undisturbed environment. We examined the fine-scale distribution and orientation of ∼230,000 organisms belonging to three groups of Rhizaria, including the mixotrophic taxa Acantharia and Collodaria, and the non-mixotrophic Phaeodaria. Overall, our results suggest that mixotrophic protists have better ability to control their position within the water column (depth, orientation) than non-mixotrophic ones. Our observations also point to several steps in the obscure life cycle of the mixotrophic Collodaria, during which *de novo* symbiont acquisition appears to involve active fine-scale buoyancy control. Taken together, these unprecedented results demonstrate that complex ecological behaviour can be achieved by “simple” organisms.

## 1 Introduction

### 1.1 Mixotrophy in marine protists

Mixotrophy – the ability to use alternate sources of nutrients – exists in plants and metazoans, but is much more common in planktonic organisms, especially protists (single-celled eukaryotes) (Schmidt et al., 2013). Because mixotrophy enables the occupation of many ecological niches, mixotrophic planktonic protists, which combine phototrophic and heterotrophic nutrition (Caron, 2016b), are ubiquitous and dominate both freshwater and marine ecosystems (Mitra et al., 2016; Stoecker et al., 2017). Planktonic protists are the most abundant eukaryotes in pelagic ecosystems and contribute significantly to global plankton biomass (Caron et al., 2017). Involved in primary production, carbon sequestration through the biological carbon pump and linking trophic levels, planktonic protists play critical ecological roles in the oceans (Ohtsuka et al., 2015; Worden et al., 2015).

### 1.2 Morphological diversity of Rhizaria

Planktonic organisms have been the subject of scientific research for two centuries (Péron and Lesueur, 1810), with a highly dichotomous view between autotrophic phytoplankton and heterotrophic zooplankton. However, many planktonic organisms actually engage in both trophic strategies but have been neglected until recently (Millette et al., 2023). First described more than a century ago (Müller, 1858; Haeckel, 1887), Rhizaria are a diverse group of protists belonging to the SAR supergroup. Most planktonic Rhizaria have a mineral skeleton, the composition of which varies between taxa: strontium sulphate in Acantharia, calcium carbonate in Foraminifera, and, opaline silica in Phaeodaria and polycystine Radiolaria (Nakamura and Suzuki, 2015; Biard, 2022). In addition, internal structures are specific to the different groups: solitary Collodaria have a large central capsule surrounded by vesicular alveoli (Anderson, 1983); Phaeodaria have a central capsule with a phaeodium (aggregate of food and waste vacuoles) (Kling and Boltovskoy, 1999). Rhizaria range in size from tens of micrometres to several millimetres (Nakamura and Suzuki, 2015; Suzuki and Not, 2015). Despite being unicellular organisms, most collodarian species exhibit colonial forms with up to tens of thousands of single cells embedded in a gelatinous cytoplasmic matrix, ranging from millimetres to metres (Suzuki and Not, 2015). Previously thought to be distinct based on physical characteristics, molecular studies have shown that solitary and colonial forms share a molecular signature, thus representing two stages of the life cycle of the same species (Biard et al., 2015).

### 1.3 Trophic ecology and photosymbiosis in Rhizaria

As no planktonic Rhizaria are cultured in the laboratory, except for a few species of Foraminifera (Kimoto, 2015), the little we know about this group comes only from *in situ* observations and sampling, whether of individuals or environmental DNA. Most Rhizaria are phagotrophic, i.e. they feed on particles, living or dead (Anderson, 1983; Biard, 2022). Some, such as Phaeodaria or benthic Foraminifera, are purely heterotrophic, feeding on suspended matter and other planktonic organisms (Kimoto, 2015; Nakamura and Suzuki, 2015). On the other hand, many epipelagic Rhizaria (e.g. Acantharia, Collodaria) are mixotrophic and host diverse photosynthetic symbionts (Decelle et al., 2015). In such an association, endosymbionts provide nutritional resources to their host and benefit in return from a favourable microenvironment, rich in nutrients and protected from predators and parasites (Yellowlees et al., 2008). However, host cells must maintain their symbionts and eventually pass them on to their offspring (Decelle et al., 2015). Endosymbiotic cells can proliferate within the host to compensate for the loss of symbionts. Most planktonic Foraminifera and Radiolaria are obligate mixotrophs: adult life stages cannot survive without their symbionts (Decelle et al., 2015).

### 1.4 Reproductive strategies of Rhizaria

Knowledge of the life cycle of Rhizaria remains scattered for most groups. Asexual reproduction by binary fission has also been reported in Foraminifera (Boudouresque, 2015; Kimoto, 2015), Phaeodaria (Nakamura and Suzuki, 2015) and Collodaria (Brandt, 1902; Anderson and Gupta, 1998). During asexual reproduction by mitosis of the host, symbionts can be transferred to the daughter cells by vertical transmission. In addition to this ability for vegetative reproduction, sexual reproduction has been reported in Foraminifera (Kimoto, 2015) and is thought to occur in other phyla of Rhizaria (Anderson, 1983; Decelle and Not, 2015; Kimoto, 2015; Nakamura and Suzuki, 2015). Indeed, the release of swarmers – small biflagellated cells (2-5 µm) – has been documented in Collodaria (both solitary and colonial, (Hollande and Cachon-Enjumet, 1953; Anderson, 1983)), Acantharia (Decelle et al., 2012), Foraminifera (Kimoto, 2015) and Phaeodaria (Kling and Boltovskoy, 1999), but whether these swarmers are gametes remains unclear (Anderson, 1983), although their morphological similarity to foraminiferan gametes has been noted (Decelle et al., 2015; Yuasa and Takahashi, 2014; Rizos et al., 2024). Finally, as sexual reproduction is extremely widespread in the eukaryotic world and was already present in the last eukaryotic common ancestor (Speijer et al., 2015), it seems reasonable to assume that different clades of Rhizaria do engage in sexual reproduction, consistent with the haploid-diploid life cycle proposed for this group (Rizos et al., 2024). However, as no vertical transmission has been reported during sexual reproduction in planktonic symbioses, it is assumed that cells produced by fertilisation acquire their symbionts *de novo* in the environment (Decelle et al., 2015), but how potential symbionts are specifically encountered and recognised remains an open question.

### 1.5 In situ imaging for advancing Rhizaria ecology

While Rhizaria have historically been overlooked due to their damage during sampling with classical plankton sampling tools (e.g. nets, pumps), recent studies based on underwater imaging – a non-destructive approach – have shed light on the ecological roles of Rhizaria (Caron, 2016a). By resolving high spatiotemporal distributions as well as the relationship between the organisms and their environment, *in situ* imaging revealed contrasted patterns in the distribution of these unicellular organisms (Biard and Ohman, 2020), but also highlighted their important contribution to the oceanic carbon biomass (Biard et al., 2016; Drago et al., 2022). *In situ* imaging can also shed light on the specific position or behaviour of organisms, such as their true occupancy volume and preferred orientation (Gaskell et al., 2019), but also on potential predation behaviour (Mars Brisbin et al., 2020).

### 1.5 Objectives of the study

As both mixotrophic and non-mixotrophic organisms coexist within Rhizaria, this group allowed us to study the ecological specificities required for mixotrophy: how does the fine-scale distribution of mixotrophic and non-mixotrophic organisms differ? Are life stages and the *de novo* acquisition of symbionts related to the position in the water column? By applying deep learning methods to automate the processing of high resolution *in situ* imaging data of planktonic Rhizaria from the NW Mediterranean, across a front in an oligotrophic environment – a habitat where Rhizaria are important (Panaïotis et al., 2024) – we (i) resolve the fine-scale distribution of these organisms, especially comparing mixotrophs and non-mixotrophs; (ii) reveal individual morphological specificities; and (iii) fill gaps in the ecological knowledge – mainly the reproductive cycle – of these mixotrophic organisms.

## 2 Materials and methods

### 2.1 Collect of plankton imaging data

Imaging data was collected using the In Situ Ichthyoplankton Imaging System (ISIIS) (Cowen and Guigand, 2008). ISIIS was deployed during the VISUFRONT cruise in July 2013 in the NW Mediterranean Sea to investigate the distribution of plankton across the Ligurian Front, a permanent front in the Ligurian Sea (NW Mediterranean). Sampling consisted of six transects performed across the front, perpendicular to the coast, for a duration of 6 to 8 hours each, during which ISIIS was deployed in a tow-yo fashion between the surface and ∼100 m. Among the six transects, three were conducted during the day and three during the night.Diel cycles were not explicitly investigated in this study. However, Faillettaz et al. (2016), working on the same dataset, found no significant differences in vertical distribution between day and night transects for Acantharia and colonial Collodaria, while solitary Collodaria were found slightly deeper during the day. ISIIS is an imaging instrument targeting planktonic organisms from 250 µm to 10 cm in size, i.e. meso- and macroplankton, where the image is acquired from the side. However, quantitative estimates are reliable only for organisms larger than ∼1 mm ESD (equivalent spherical diameter) (Panaïotis et al., 2022), meaning that smaller organisms may be underrepresented in the dataset. It features a large field of view (10.5×10.5×50 cm, about 5,500 cm^3^) which not only provides a very high sampling rate (> 100 L s^-1^), but also ensures that the centre of the imaged volume and organisms herein are barely disturbed by the presence of the instrument. In addition, the shadowgraphy technique used in ISIIS is particularly well suited to imaging transparent planktonic organisms and their internal structure. However, organisms may be horizontally distorted depending on the towing speed and the acquisition rate of the ISIIS line scanning camera. In our case, the organisms were laterally compressed so that a circle appears as a vertical ellipse and the organisms were therefore imaged smaller than in reality. This deformation was estimated by computing the ratio of height to width of organisms assumed to be spherical, and the width was corrected accordingly when evaluating the size of the organisms. Furthermore, due to the oscillating sampling pattern, ISIIS has a pitch (the angle of the instrument relative to the horizontal) that is recorded to allow for later correction. Finally, ISIIS also continuously records environmental data: temperature, salinity, fluorescence and oxygen.

### 2.2 Detection and classification of planktonic organisms

ISIIS recorded almost 44 hours of data (equivalent to a 185 million pixel-long image), the processing of which had to be automated. The first processing step was to detect planktonic organisms in the raw images. Using a content-aware segmentation pipeline based on a convolutional neural network (CNN) (Panaïotis et al., 2022), more than 20 million potential planktonic organisms were extracted. In a second step, these organisms were automatically sorted into 24 taxonomic groups using a CNN classifier (MobileNetV2) previously trained and evaluated on ISIIS data (Panaïotis et al., 2025). The 1.8 million organisms sorted within Rhizaria were selected for a finer classification step into 14 categories, using a specifically trained classification model (Figure 1). For both classifiers, data augmentation included image flipping, so that specific orientations are unlikely to be the result of a classification artefact. The identification of the 1.8 M images could not be manually validated. Instead, model performance was estimated on an independent dataset and uncertain predictions – below a probability threshold calculated so that 90% of errors occurred below this threshold – were discarded. This method decreased recall but improved precision, i.e. concentrations are underestimated but distribution patterns are preserved (Faillettaz et al., 2016).

**Figure 1.**
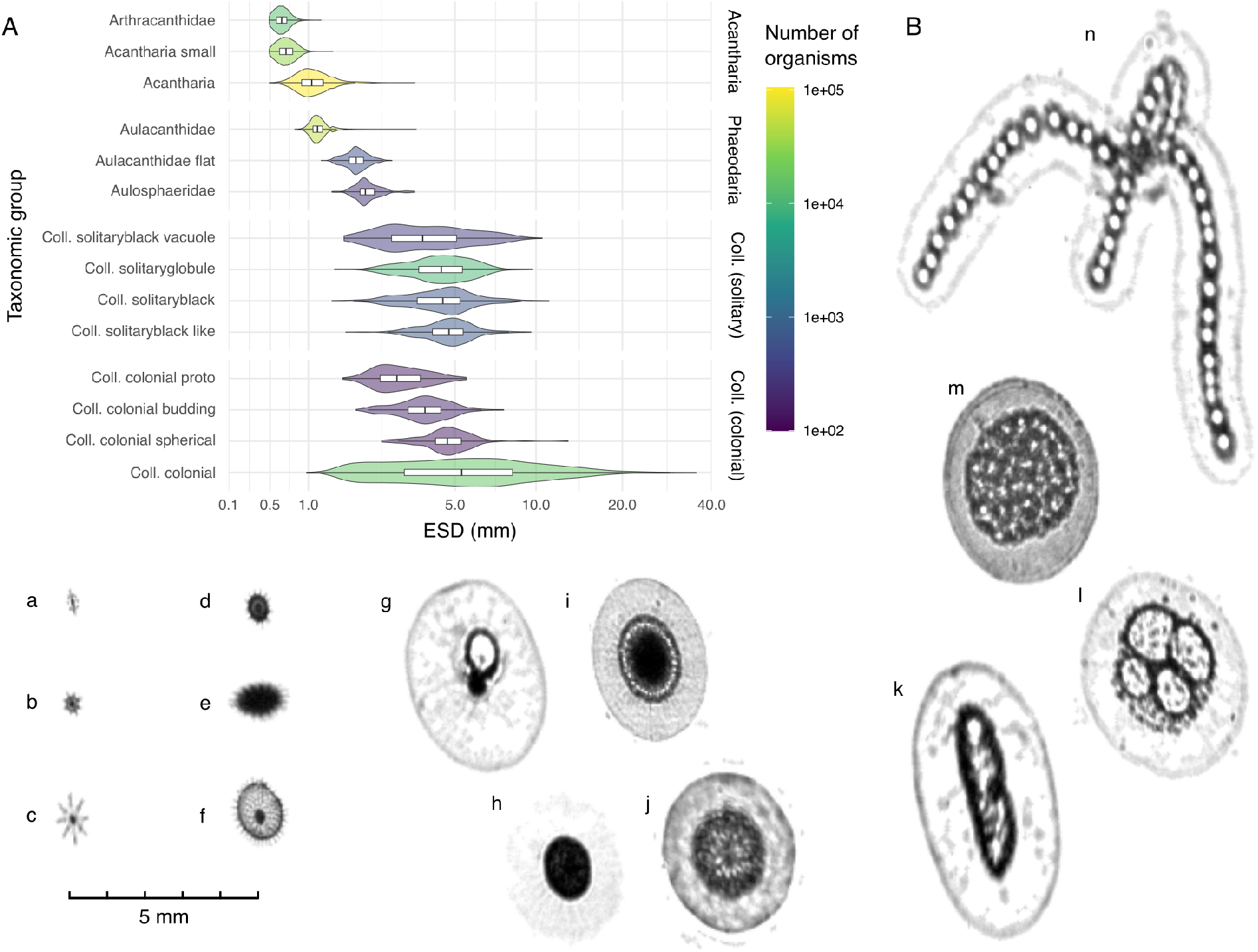
Composition of the dataset. **(A)** Equivalent Spherical Diameter (ESD) distribution and number of organisms per taxonomic group (Coll. = Collodaria). **(B)** Examples of ISIIS images for each taxonomic group: (a) Arthracanthida, (b) Acantharia small, (c) Acantharia other, (d) Aulacanthidae, (e) flat Aulacanthidae, (f) Aulosphaeridae, (g) Collodaria solitary with vacuole, (h) Collodaria solitaryglobule, (i) Collodaria solitaryblack, (j) Collodaria solitaryblack like, (k) Collodaria budding, (l) Collodaria proto colony, (m, n) Collodaria colonial.

### 2.3 Morphological feature extraction and additional annotations

After classification, each organism was characterised morphologically by measuring a set of features, including proxies for size (e.g. area, perimeter), transparency (e.g. mean grey level) and *in situ* position (e.g. orientation of the major axis). Each element of the dataset consisted of an image, a taxonomic label, a set of features and associated environmental data, from which concentrations and vertical distributions were computed. In addition, the positions of specific structures of three categories were manually recorded using a keypoint annotation tool^1^. Approximately 350 solitary Collodaria (∼3.5% of all solitary Collodaria) showed asymmetric vacuoles. By recording the position of both the centre of the nucleus and the tip of the largest vacuole, vacuole orientation and size were computed for each organism. Similarly, the position of the phaeodium in relation to the centre of the organism was recorded for 224 Aulosphaeridae (∼0.5% of all Aulosphaeridae). Finally, the orientation of Arthracanthida (Acantharia) was assessed manually by recording the position of the extremities of the longest spicule in 232 organisms (∼4.5% of all Arthracanthida). All orientation values were corrected for the pitch of ISIIS.

### 2.4 Environmental data processing

Abnormal environmental values (e.g. negative temperature and oxygen) were first removed. Density was computed from temperature and salinity. Finally, a bilinear interpolation of the transects was performed using distance from shore in *x* (200 m steps) and depth in *y* (0.5 m steps) to obtain complete images of the transects. The deep chlorophyll maximum (DCM) depth was computed from the interpolated data as the depth of maximum fluorescence. As it highlighted well the downwelling of oxygenated waters, oxygen and Rhizaria concentration values along the DCM were extracted to investigate whether Rhizaria distribution was affected by these vertical water movements.

All analyses were performed using `R` version 4.1.2. Data processing and interpolation were performed using the `dplyr` and `akima` packages respectively. Plots were generated with `ggplot2` using the colour blind friendly `viridis` and `cmocean` colour scales.

## 3 Results

### 3.1 A comprehensive dataset

Among the ∼8 million planktonic organisms imaged with ISIIS, approximately 230,000 were identified as Rhizaria and classified into 14 taxonomic and/or morphological categories, belonging to three larger groups: Acantharia, Phaeodaria, and Collodaria (for which we distinguished colonial and solitary stages). Each morphological category contained between 147 and 104,455 individuals, covering a size range from 0.4 mm (ESD) for Acantharia to 35 mm for colonial Collodaria (Figure 1). Acantharia (n ≈ 150,000) dominated the dataset. Within Acantharia, the categories Acantharia small and Acantharia othe’ are primarily distinguished by size and likely encompass multiple species. Aulacanthidae (n ≈ 50,000; Phaeodaria) were also abundant. In the NW Mediterranean, the most common Aulacanthidae species is *Aulacantha scolymantha*, which is spherical in shape and possesses long siliceous spines (Nakamura and Suzuki, 2015). The organisms imaged here are consistent with this morphology, with an ESD generally around 1 mm or slightly larger; indicating that smaller individuals may have been missed due to the limited pixel resolution of ISIIS. Collodaria were less abundant (n ≈ 10,000 solitary cells; n ≈ 15,000 colonies).

Organisms were distributed in a strongly stratified water column (Figure S1), with a thermocline/pycnocline at about 10 m depth. The front was well marked, delimiting fresher water inshore from saltier water offshore. The DCM was located between 75 m offshore and 50 m inshore where it was more spread out. Finally, oxygenated waters were found between the thermocline and the DCM, highlighting two tongues of downwelling waters that were also visible in the temperature. Overall, these features are typical of the oligotrophic summer period in the Ligurian Sea.

### 3.2 Vertical distribution of Collodaria depended on life stages

Almost all known Collodaria are mixotrophic (Biard, 2022; Nakamura et al., 2023) and do not all possess the silicified skeleton typical of polycystine Radiolaria. Beyond the typical solitary and colonial forms, additional forms were detected by ISIIS. These forms could correspond to the transition phase (i.e. budding – being a continuous succession of binary fissions) between the solitary and colonial stages (Figure 2D). Overall, solitary forms were found close to the DCM, with the exception of vacuole-bearing organisms found deeper below the DCM; while colonies were more dispersed in the water column although still centred on the DCM (Figure 2C). Finally, the vacuoles of the 350 manually annotated solitary organisms showed a clear orientation towards the surface (Figure 2B) and the size of the vacuoles increased with depth (Figure 2A).

**Figure 2.**
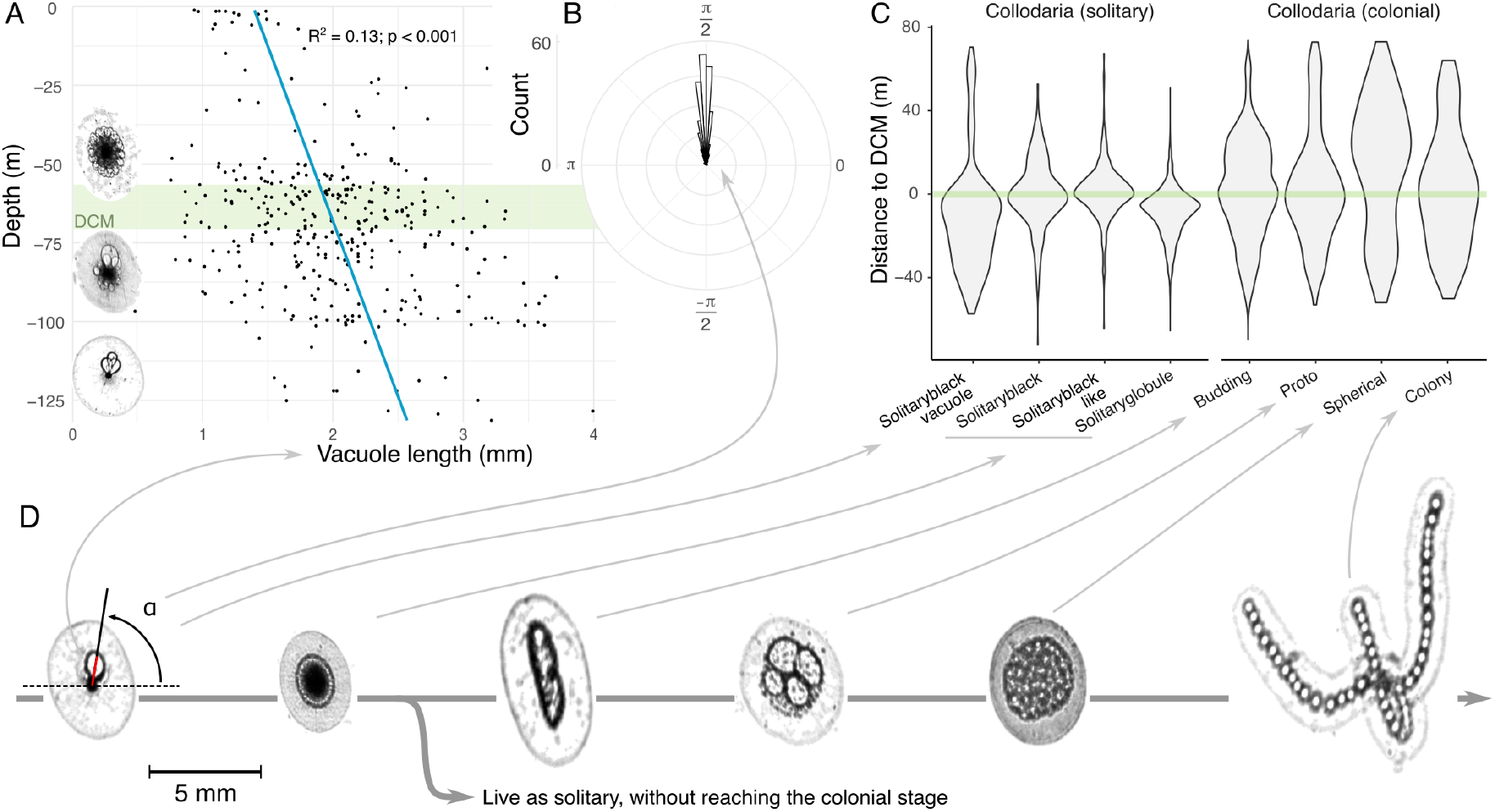
Vertical distribution of Collodaria and vacuole characteristics. **(A)** Vacuole length (shown as a red line in **D**) VS depth in solitary Collodaria. The green band represents the DCM. Typical organisms from three depths are shown on the left of the plot. **(B)** Angle of vacuole position (shown in **D**) in solitary Collodaria. **(C)** Distribution relative to DCM for group of Collodaria. **(D)** Morphology of Collodaria detected by ISIIS and their putative chronological order during the life cycle of Collodaria.

### 3.3 Acantharia had disparate vertical distributions

Three subgroups of Acantharia, smaller overall than Collodaria but also exclusively bearing a strontium sulphate skeleton, were identified and showed distinct vertical distributions (Figure 3A): Arthracanthida were found around the DCM, other Acantharia were found well above the DCM, just below the surface, while small Acantharia showed an intermediate distribution between these, with some organisms being close to the surface and others were around the DCM, suggesting a mix of different subgroups. Moreover, Arthracanthida displayed a preferential *in situ* orientation with their largest spicule oriented vertically (Figure 3BC).

**Figure 3.**
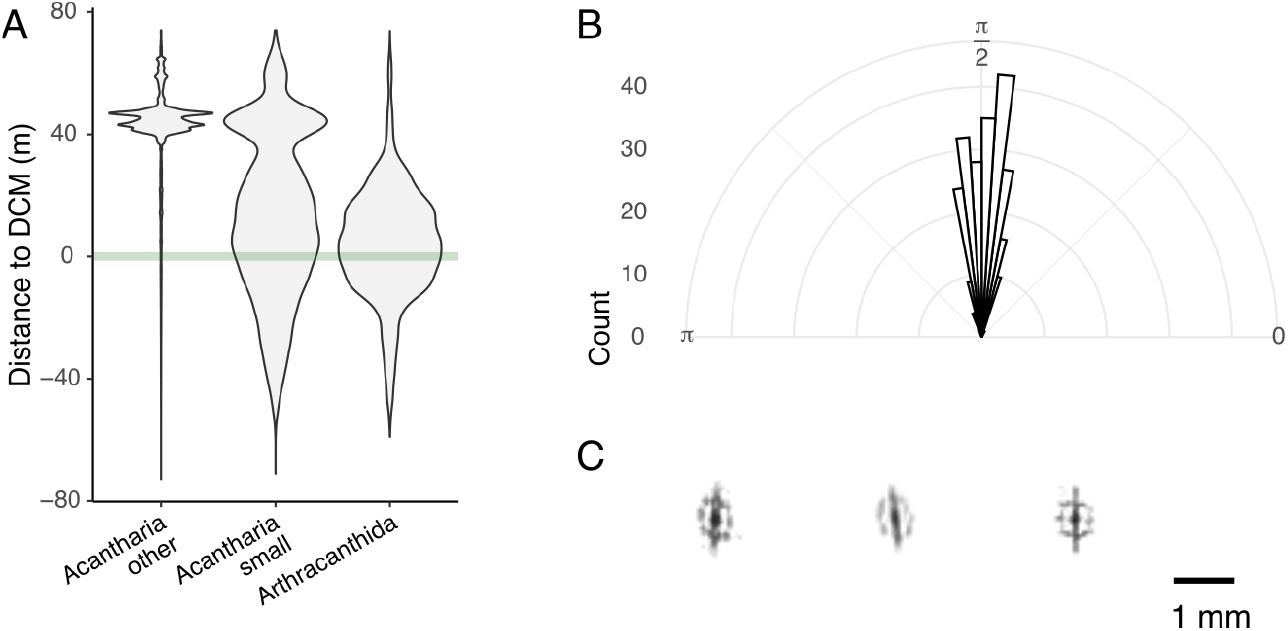
Vertical distribution of Acantharia and natural orientation in the water. **(A)** Distance to DCM for Acantharia groups. **(B)** *In situ* orientation of Arthracanthida. **(C)** Examples of Arthracanthida images.

### 3.4 Phaeodaria distribution differed from mixotrophic Rhizaria

Unlike Collodaria and Acantharia, Phaeodaria lack photosymbionts and are strictly heterotrophic. The two subgroups of Aulacanthidae were found around the DCM (Figure 4C), but appear to be moved downwards by sinking waters: when oxygen concentration is higher on the DCM (green line in Figure 4A), corresponding to sinking water, the concentration of Aulacanthidae is lower (Figure 4AB, S2). This was not the case for other mixotrophic Rhizaria dwelling around the DCM (Figure S3). Some organisms identified as Aulacanthidae had an ellipsoidal shape and were larger than the typical spherical Aulacanthidae. These organisms, termed “Aulacanthidae flat”, could not be identified with certainty but were assigned to the same family based on their identical vertical distribution to typical Aulacanthidae (Figure S2), and were all horizontally oriented (Figure 4D). Aulosphaeridae were typically found deeper, below the DCM (Figure 4C). In these organisms, the phaeodium was found to be typically positioned towards the bottom of the cell (Figure 4E).

**Figure 4.**
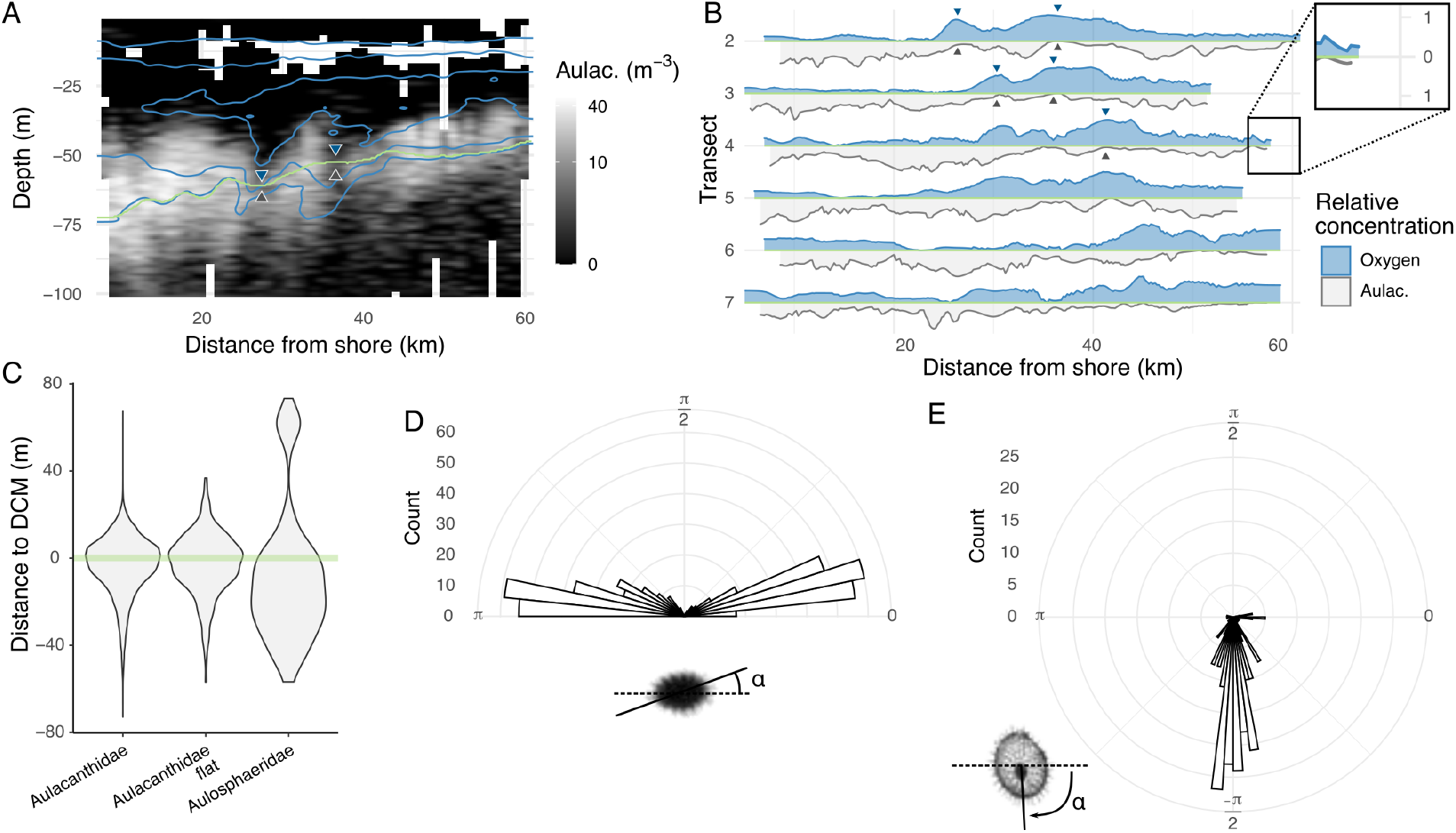
Distribution and orientation of Phaeodaria. **(A)** Distribution of Aulacanthidae along a transect. White lines are the 210, 230 and 250 µmol kg^-1^ oxygen isolines highlighting downwelling waters around 28 and 37 km offshore. The green line is the DCM. Note that the colour scale is log-transformed. **(B)** Relative Aulacanthidae and oxygen concentration along the DCM (green line in **A**). Blue arrowheads highlight the downwelling of high oxygenated waters corresponding to lower concentrations of Aulacanthidae, highlighted by grey arrowheads. **(C)** Distance to DCM for Phaeodoaria groups. **(D)** *In situ* orientation of flat Aulacanthidae. **(E)** Angle of phaeodium position in Aulosphaeridae.

## 4 Discussion

### 4.1 Mixotrophy in giant protists

#### 4.1.1 Collodaria: a complex life cycle constrained by mixotrophy

We observed Collodaria organisms with diverse morphologies and different vertical distributions. In particular, some solitary cells exhibited vacuoles, systematically oriented towards the surface, with their size decreasing with decreasing depth. Here, we try to relate these observations to the current knowledge of the poorly described life cycle of Collodaria, in relation to the environment. In particular, we seek to understand how *de novo* symbiont acquisition can occur in unicellular organisms that lack a flagellum and are unable to swim, in the absence of vertical symbiont transmission.

Although Collodaria are typically found in the epipelagic layer because they host symbionts, buoyancy loss and morphological changes occur concurrently with swarmers release (Anderson, 1983). Although the exact location of swarmer release is not known, as their fate, there is indirect evidence that a hypothetical fertilisation might occur at depth. First, fertilisation is thought to occur at depth in both Acantharia and Foraminifera (Decelle et al., 2013; Kimoto, 2015). Second, Collodaria has previously been detected at depth by metabarcoding (Pernice et al., 2016; Faure et al., 2019), while almost all known species host photosymbionts (Nakamura et al., 2023) and therefore typically reside in the photic zone. More specifically, these molecular signals are identical to those of the clades detected in the epipelagic zone, highlighting the presence of the same species both at the surface and at depth. Collodaria was also detected in the picoplankton (0.2-2 µm, (Mucko et al., 2018)), while all Collodaria are typically larger than 1 mm. These elements, very small and detected at depth, are likely to correspond to swarmers. Finally, Collodaria swarmers have been shown to contain a crystal of strontium sulphate in their cytoplasm (absent in the juvenile stages), which may act as a ballast to facilitate the swarmer’s descent to depth (Yuasa and Takahashi, 2014).

If fertilisation does occur at depth, the newly formed organism then has to reach the photic zone where adults live. This change in depth is likely facilitated by the vacuoles, which are thought to contain lipids (Anderson, 1983; Febvre-Chevalier and Febvre, 1994). All the larger vacuoles we detected in solitary Collodaria were oriented upwards, suggesting that they provide positive buoyancy to the cell. The decreasing size of vacuoles as they approach the surface, combined with the reorganisation of vacuoles around the nucleus (Figure 2A), is consistent with a shift from positive buoyancy to stability as organisms reach the target habitat. Thus, we hypothesise that vacuole-bearing cells could correspond to newly formed organisms, migrating from the fertilisation site to their next habitat: the DCM, a source of food and potential symbionts. These unicellular organisms therefore appear to be able to actively control their buoyancy in order to meet their energy requirements for mixotrophy.

Several lines of evidence support the absence of vertical transmission of symbionts: symbionts are lost before gametogenesis and swarmers are too small to host symbionts (Decelle and Not, 2015). This suggests a need for the *de novo* acquisition of symbionts from the environment, which in the oligotrophic waters of the Mediterranean Sea during summer, are only abundant in the DCM, where Collodaria could also feed on various planktonic organisms (Biard et al., 2015).

With these acquired energetic resources, a solitary cell can enter in a budding phase (i.e. a sequence of binary fissions) to form a new colony by vegetative reproduction (Hollande and Cachon-Enjumet, 1953), which continues to grow by cell division (Anderson and Gupta, 1998), consistent with the colonial cycle proposed for Radiolaria (Yuasa and Takahashi, 2014; Rizos et al., 2024). Meanwhile, symbionts are able to reproduce within the host (Decelle et al., 2015), so that, at some point, the host does not need to acquire new symbionts. Our data revealed that colonies are more dispersed in the water column than solitary cells, in line with previous observations (Faillettaz et al., 2016). This could be due to a weaker ability to control buoyancy, or the result of less habitat pressure to stay close to the DCM, allowing the use of a larger habitat. It is also possible that colonies alternate between habitats: more nutrients at depth and maximum sunlight exposure of symbionts closer to the surface.

Beyond the hypothesis developed above to explain our observations, in particular the vacuoles, we propose alternative hypotheses. A first one could be that lipid vacuoles are used for ascent after fertilisation, without a specific target such as the DCM, and that only cells that reach the DCM (and stop there) survive. This hypothesis would explain the presence of few vacuole-bearing solitary Collodaria very close to the surface. However, this strategy seems sub-optimal, as it induces higher mortality rates than our main hypothesis, and should therefore not have been maintained by natural selection. Another, less likely, hypothesis is that our plankton detection and classification algorithms are biased towards upward-facing vacuoles, ignoring other orientations. However, these organisms are quite large and are detected using a naive grey thresholding of the images, so there is no reason for a bias to occur at the detection stage. We then inspected ∼300 solitary Collodaria prior to automated classification and all vacuoles detected were up-facing, making this alternative hypothesis unlikely.

Overall, although we cannot prove that the above steps constitute the life cycle of Collodaria, our observations tell a coherent story. To confirm this, vacuole-bearing organisms would need to be sampled *in situ* to check for the absence of symbionts, as this stage should occur prior to symbiont acquisition. However, this does not seem feasible given the very small number of observations: Collodaria are generally scarce, and only ∼350 solitary Collodaria with vacuoles were detected in the 17×10^6^ L sampled in 44 hours. Moreover, the low proportion of vacuole-bearing cells compared to the total number of solitary Collodaria (∼10,000) suggests that the duration of the vacuole stage would be very short or rare compared to the time spent in the DCM. Another solution could be to use multispectral *in situ* imaging to detect fluorescence within organisms (Franks and Jaffe, 2001; Zawada, 2003; Liu et al., 2018), but again this, would require a very high sampling rate, of the same order of magnitude as ISIIS (> 100 L s^-1^).

#### 4.1.2 Acantharia: various mixotrophic strategies

Photosymbiont-hosting Rhizaria are expected to be found near the surface for a maximum exposure to sunlight, but this environment is particularly nutrient poor, hence the importance of mixotrophy. However, the two mixotrophic groups we studied (Acantharia and Collodaria) had very different distributions. In Acantharia, three distinct vertical patterns were detected (Figure 3). However, the limited pixel resolution of ISIIS (1 px = 51 µm) prevented a finer identification, which could have explained the bimodal distribution of small Acantharia. Larger, probably symbiont-bearing Acantharia were found very close to the surface, in agreement with previous findings (Michaels, 1988). Collodaria – also mixotrophic – had a very different distribution from Acantharia: solitary cells were found around the DCM while colonies were more dispersed (although colonies can accumulate close to the surface; (Suzuki and Not, 2015)). One hypothesis to explain this difference in strategies is that Acantharia-Phaeocystis symbioses produce high amounts of dimethylated sulphur compounds, which are likely to act as antioxidants, enabling the holobiont to cope with oxidant products generated by photosynthesis and high irradiance exposure (Decelle et al., 2012). The differences in vertical distribution between Acantharia subgroups could also reflect differences in their symbiont communities. Indeed, different Acantharia taxa are known to host different photosynthetic symbionts, with distinct light requirements (Decelle et al., 2015). Similarly, the difference in distribution between Acantharia and Collodaria could partly reflect the different nature of their symbionts, Acantharia hosting mainly Phaeocystis while Collodaria host dinoflagellates (Decelle et al., 2015).

### 4.2 Unicellular organisms able to control their position

#### 4.2.1 Vertical position

Collodaria, Aulacanthidae and Arthracanthida had a vertical distribution centred around the DCM – a potentially favourable environment (for feeding, symbiont acquisition) – which is likely an indicator of an active buoyancy control. Such active buoyancy control has been reported previously in Rhizaria (Anderson, 1983; Michaels, 1988; Febvre-Chevalier and Febvre, 1994; Jon Furbish and Arnold, 1997). The vacuoles of solitary Collodaria – systematically oriented towards the surface – and the relationship between vacuole size and depth mentioned above are further elements in favour of active buoyancy regulation.

Although it was initially thought that non-motile unicellular planktonic organisms float freely in the water column (Basterretxea et al., 2020), several buoyancy control mechanisms have been discovered, mostly consisting of accumulation of substances that modify cell density (e.g. lipid droplets in Acantharia, polycystine Radiolaria and Phaeodaria, (Febvre-Chevalier and Febvre, 1994)) or active modification of the cell shape and volume (e.g. myoneme contraction in Acantharia, (Febvre, 1981)), but see Febvre-Chevalier and Febvre (1994) for a review. Beyond protists, cyanobacteria such as *Trichodesmium* can regulate their buoyancy through gas vacuoles (Walsby, 1978).

The DCM is typically located around the density gradient (Figure S1) that separates the nutrient-poor surface layer from a light-limited deep layer (Herbland and Voituriez, 1979), thus corresponding to favourable conditions for phytoplankton growth. However, neutral buoyancy and accumulation on density gradients of phytoplankton cells can also contribute to the formation of DCMs (Lofton et al., 2020). Therefore, a passive accumulation on the same density gradient of other organisms also explains their distribution. This is likely to be the case for Aulacanthidae (Phaeodaria), whose distribution appeared to be affected by small-scale downwellings, which could be a sign of very limited buoyancy control. Nevertheless, Aulacanthidae are thought to maintain their buoyancy thanks to a low carbon and biogenic silica density (their test is porous) (Laget et al., 2023), and Coelodendrid (Phaeodaria) can maintain neutral buoyancy *in vitro* (Swanberg et al., 1986). However, to our knowledge, no active buoyancy regulation mechanisms have been reported in the literature for Phaeodaria, in contrast to other Rhizaria groups, although some behavioural observations suggest it may occur (Nakamura et al., 2018).

#### 4.2.2 Preferred orientation

The use of *in situ* imaging with images taken from the side allowed us to resolve the *in situ* orientation of several organisms. We detected preferential orientation both from the shape of the organisms (oblate or prolate ellipsoid) and from the specific position of internal asymmetric structures in spherical organisms (e.g. lipid vacuoles, phaeodium). Positively buoyant lipid vacuoles were oriented towards the top of the spherical solitary Collodaria cells; whereas the phaeodium, a denser aggregate of waste and food (Kling and Boltovskoy, 1999), was located towards the bottom of the also spherical Phaeodaria cells. A similar preferential orientation has previously been reported *in situ* in Foraminifera with bubble capsules positioned towards the surface (Gaskell et al., 2019).

Similar to vertical position, cell orientation in unicellular planktonic organisms was considered to be random due to small-scale turbulence (Font-Muñoz et al., 2019). However, cell orientation conditions several essential functions such as reproduction, sensing, metabolism or locomotion, and non-uniform orientation distributions are the norm (Basterretxea et al., 2020). In non-motile unicellular organisms with homogeneous density, cell orientation is mediated by fluid-cell interactions at the microscale, such that cells adopt a hydrodynamically favourable orientation (Basterretxea et al., 2020). Alternatively, orientation may be mediated by differences in the density of internal structures.

Several studies have focused on the orientation of different particles, revealing a preferential horizontal orientation in regions of low shear (Nayak et al., 2018). Furthermore, the time spent horizontally increased with the aspect ratio of the particles, in line with the predictions of Jeffery’s theoretical model (Jeffery, 1922). Similar results have been obtained for diatom chains *in situ* (Malkiel et al., 1999; Talapatra et al., 2013; McFarland et al., 2020) and ex situ (Karp-Boss et al., 2000). In addition, the horizontal orientation of phytoplankton colonies appears to be ecologically beneficial as it increases the area exposed to sunlight, which could lead to an increase in photosynthetic activity (Bricaud and Morel, 1986; McFarland et al., 2020). In contrast, pennate diatoms were found to vertically reorient when sinking from surface turbulent waters (Font-Muñoz et al., 2019).

The above literature shows that a horizontal orientation is expected for non-motile plankton. This was the case for our oblate, flat Aulacanthidae. Nonetheless, Arthracanthida (Acantharia) showed a different preferred orientation, with the two longest and thickest spicules in the vertical direction. Since their skeleton consists of strontium sulphate, the densest oceanic biomineral known (3.96×10^3^ kg m^-3^) (Marszalek, 1982), their vertical orientation could result from a passive equilibrium imposed by the weight of the skeleton, although the presence of an internal structure of higher or lower density influencing cell orientation cannot be excluded. In addition, Acantharia have been reported to actively deploy long cytoplasmic extensions that could be involved in predation by capturing food particles, although this is not certain (Mars Brisbin et al., 2020). These structures may also be involved in other functions, including buoyancy control.

Overall, our results seem to indicate that, at least within the studied taxa, mixotrophic protists show a better ability to actively control their buoyancy and position within the water column than non-mixotrophic ones, although this pattern may not be universal (Nakamura et al., 2018). Ultimately, our results challenge the historical view of a completely passive life for planktonic protists, and even more so for the mixotrophic ones.

### 4.3 New insights on mixotrophy

Thus, our results provide new insights into the ecology of mixotrophic protists, by suggesting how symbiont acquisition may be mediated within the life cycle. We also show that mixotrophic protists can exploit different ecological niches.

In this study, we highlight that the transition between pure heterotrophy and mixotrophy could shape the life cycle and distribution of Collodaria, in which fine buoyancy control enables the *de novo* acquisition of symbionts. This work also supports the idea that mixotrophy enables the exploitation of habitat otherwise unfavourable for photosynthesis, albeit with different trade-offs in distribution in Acantharia and Collodaria. Finally, we also point out unexpected and often neglected behaviour of unicellular organisms. More than 170 years after the first description of “yellow cells” inside colonial Collodaria (Huxley, 1851) and in other polycystine Radiolaria (Müller, 1858; Haeckel, 1887), later identified as symbionts (Brandt, 1902), there is still a lot of unknown around these organisms. Planktonic symbioses have received much less attention than others, although they play critical ecological roles in the oceans. Mixotrophy has appeared several times during evolution, in particular within several groups of eukaryotes (Stoecker et al., 2009), making it a derived trait. Within Rhizaria, mixotrophy has appeared several times too and in different ways (e.g. endosymbiosis vs. kleptoplastidy, “red” vs. “green” symbionts, (Stoecker et al., 2009)). Unsurprisingly, this trait appears to influence the habitat use: in the absence of vertical transmission, mixotrophic planktonic protists are subject to a balance of selection pressure between foraging for food or symbionts. In heterotrophic organisms, however, selection pressure is different. For Phaeodaria, which feed on particles in suspension, a limited ability to regulate buoyancy could be consistent with a more passive foraging strategy, although this remains speculative given the limited number of groups studied here and the existence of active buoyancy control in some heterotrophic protists (Nakamura et al., 2018).

In the continuity with previous work using *in situ* imaging to reveal preferential orientation (Gaskell et al., 2019) or potential predation behaviour (Mars Brisbin et al., 2020) in protists, this work demonstrates how high-throughput *in situ* imaging, combined with deep learning-based classification, provides a powerful and non-destructive approach to uncover the poorly understood ecology of fragile planktonic protists, including their distribution, life cycle, and posture in the water column.

## Data availability

Data supporting this study is available online (https://www.seanoe.org/data/00851/96266/) (Panaïotis et al., 2023), as well as the code to reproduce the analyses (https://zenodo.org/records/22301619) (Panaïotis, 2026).

## Acknowledgement

The authors acknowledge officers and crew of the R/V Tethys 2, as well as the additional scientists who took part in the cruise: F Lombard and M Lilley. This study is part of project “World Wide Web of Plankton Image Curation”, funded by the Belmont Forum through the Agence Nationale de la Recherche ANR-18-BELM-0003-01 and the National Science Foundation (NSF) ICER1927710. Data acquisition during the VISUFRONT cruise was funded by the Partner University Fund and supported by the French Oceanographic Fleet through ship time. This work was granted access to the HPC resources of IDRIS under the allocation 2021-AD011013092 made by GENCI. We are also grateful to the Roscoff Bioinformatics platform ABiMS (http://abims.sb-roscoff.fr), part of the Institut Français de Bioinformatique (ANR-11-INBS-0013) and BioGenouest network, for providing computing resources. TP’s doctoral fellowship was granted by the French Ministry of Higher Education, Research and Innovation (contract 3500/2019).

1 https://github.com/luiscarlosgph/keypoint-annotation-tool

