## Supplementary for "High throughput *in situ* imaging reveals complex ecological behaviour of giant marine mixotrophic protists"

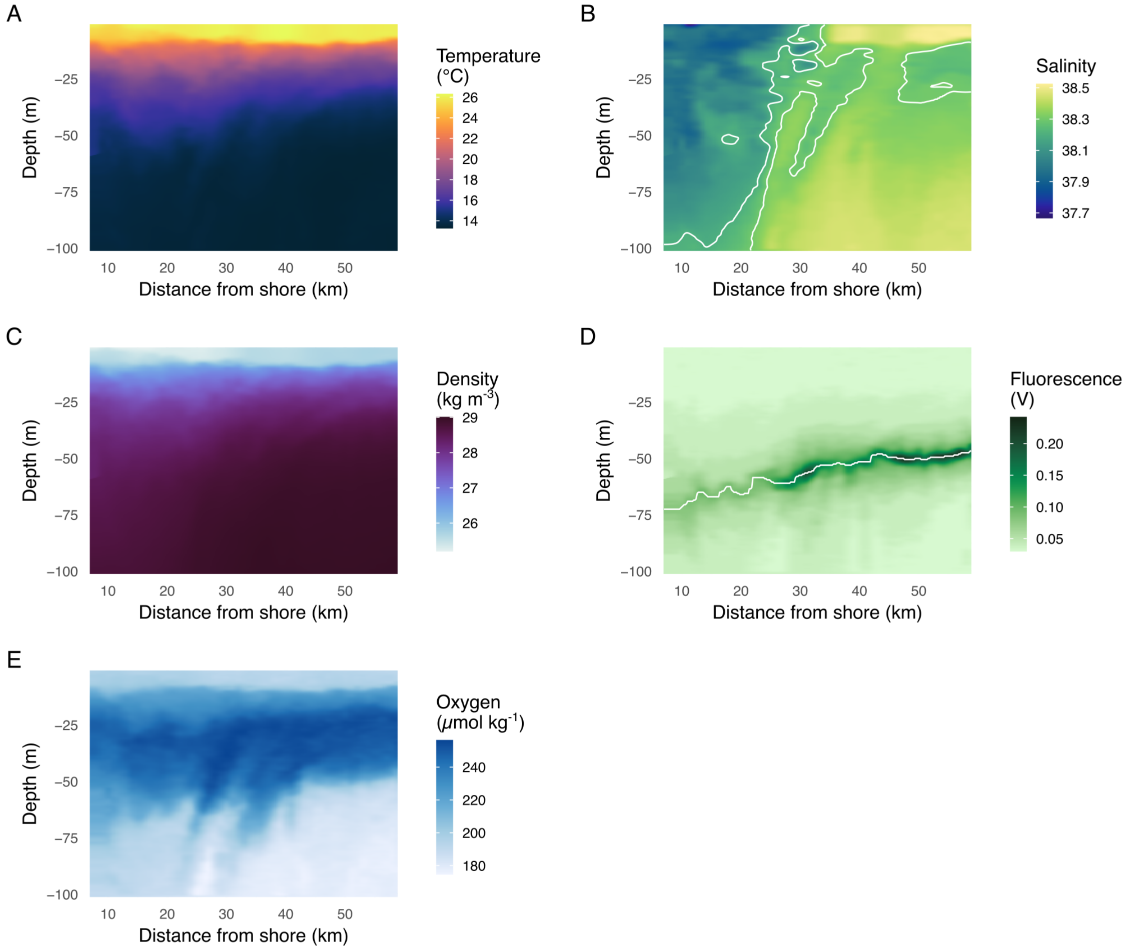


Figure S1: Environmental data along one transect, representative of the other transects. **(A)** temperature, **(B)** salinity with 38.2 and 38.3 isohalines delimiting the front, **(C)** density anomaly, **(D)** fluorescence with DCM represented as a white line, **(E)** oxygen.

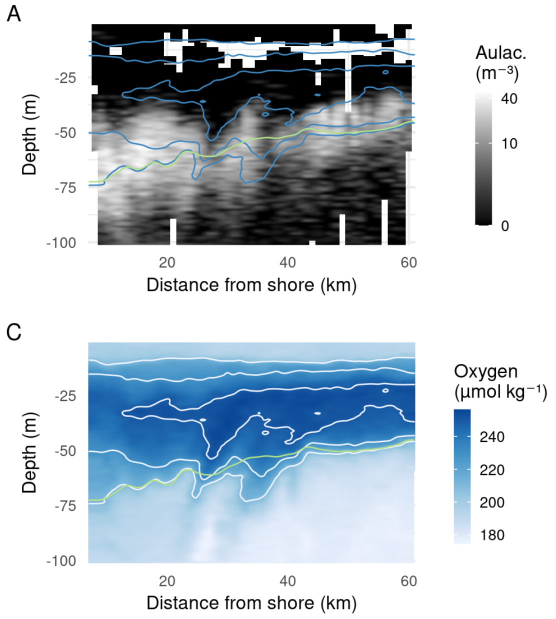


Figure S2: Effect of downwelling waters on Aulacanthidae distribution along a transect, representative of the other transects. Distributions of **(A)** Aulacanthidae, **(B)** Flat Aulacanthidae and **(C)** oxygen concentration. Blue **(A, B)** and white **(C)** lines represent oxygen isolines (210, 230 and 250 µmol $kg^{-1}$), the green line highlights the DCM.

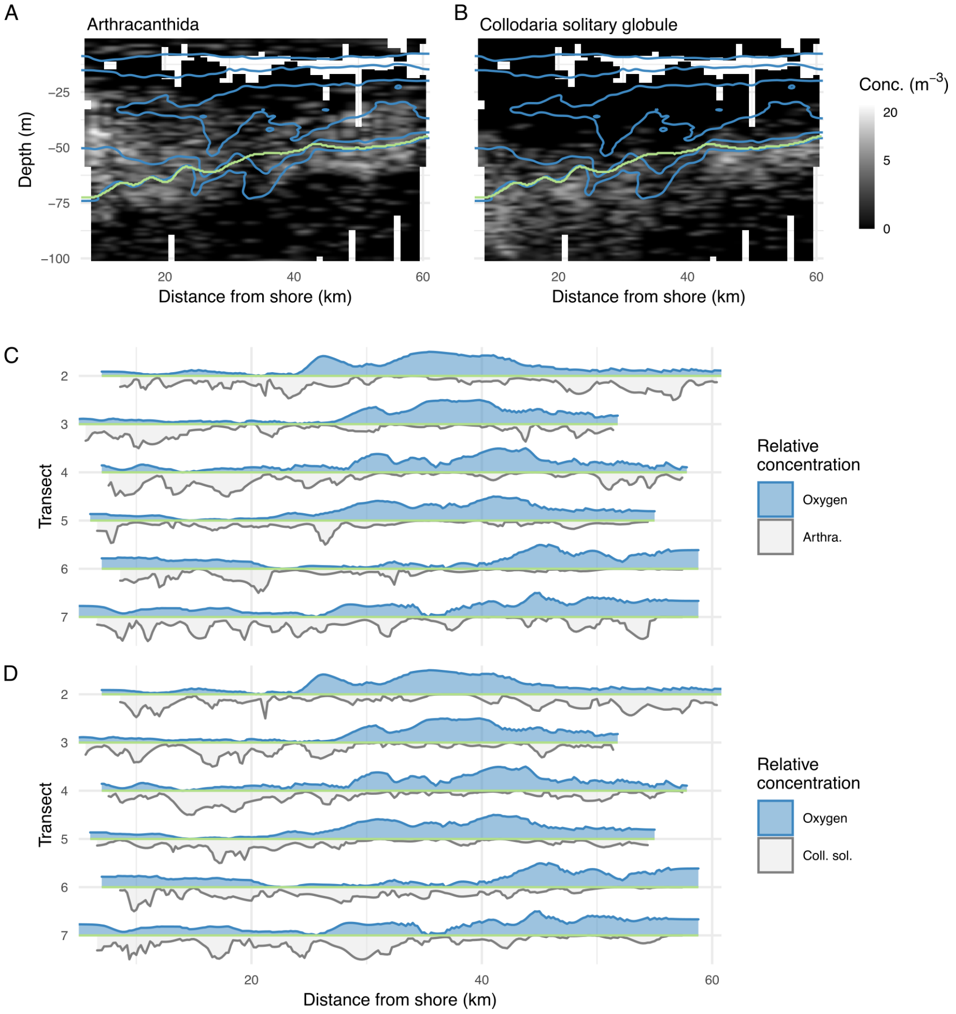


Figure S3: Absence of effect of downwelling waters on other Rhizaria. Distributions of **(A)** Arthracanthida and **(B)** solitaryglobule Collodaria along a transect. Blue lines are the 210, 230 and 250 µmol kg$^{-1}$ oxygen isolines highlighting downwelling waters around 28 and 37 km offshore. The green line represents the DCM. Note that the colour scale is log-transformed. Relative **(C)** Arthracanthida, **(D)** Collodaria solitary globule (grey) and oxygen (blue) concentration along the DCM (green line in **(A, B)**), in the [0, 1] range.
